# Sustained Volumetric Compression Induces Cell Jamming and Primes Breast Cancer Cells for Enhanced Post-Compression Migration and Invasion

**DOI:** 10.64898/2026.08.08.743678

**Authors:** Marjan Ghanbariabdolmaleki, Justin Caron, Anpreet Dhaliwal, Gabriel Medina, Dominic Mak, Raina Prasad, Jordan Ziesse, Shengjie (Patrick) Zhai, Shue Wang

## Abstract

During tumor growth and progression, cancer cells are exposed to sustained physical confinement and volumetric compression that can alter cell volume, cytoskeletal organization, mechanotransduction, and invasive behavior. However, whether breast cancer cells retain a compression-induced mechanical memory after release from sustained volumetric compression, and how this memory influences subsequent migration and invasion, remains poorly understood. Here, by controlling cell volume using PEG - mediated volumetric compression, we investigated the compression and post-compression recovery responses of MCF-7 breast cancer cells. Cells were compressed for four days, followed by four days of recovery after PEG removal, and analyzed using daily morphological tracking, single-cell time-lapse imaging, F-actin and YAP staining, wound healing assays, and 3D spheroid invasion assays. We show that sustained volumetric compression shifts MCF-7 cells into a compact, jammed-like, low-motility state characterized by reduced morphodynamic remodeling, suppressed collective migration, and limited spheroid invasion. In contrast, post-compression recovery induces a distinct mechanobiological state marked by increased cell area and perimeter, altered single-cell trajectories, heterogeneous F-actin remodeling, enhanced YAP nuclear localization in enlarged recovered cells, accelerated wound closure, and increased spheroid invasion and cell dissemination. These findings suggest that prior volumetric compression can prime breast cancer cells for enhanced migration and invasion after stress release, supporting post-compression recovery as a form of mechanical memory that may contribute to tumor dissemination.

## Introduction

Cancer is one of the leading causes of death in the United States and worldwide, with solid tumors accounting for the majority of cancer-related deaths due to their tendency for local invasion and distant metastasis [1, 2]. Cancer cell progression is governed by different microenvironment cues, including ECM architecture, biochemical cues, and biophysical cues, including stiffness and spatial confinement [3–5]. In vivo, tumor cells experience strong geometric constraints imposed by native tissue structures such as ducts, vessels, and stromal boundaries, which critically shape cellular organization and invasion behavior [6]. Breast cancer progression is regulated not only by biochemical cues, but also by biophysical cues within tumor microenvironment. As breast tumor grows within confined tissue spaces, cancer cells, stromal cells, ECM deposition, and surrounding normal tissue collectively generate abnormal stresses [7]. In addition, physical forces stem from both the fluid and the solid phase of a tumor, where the fluid phase corresponding to hydrostatic fluid pressure of tumor interstitial space, the ostmotic pressure owing to the transport of positive and negative ions and the existence of fixed charges in the tumor microenvironment [8, 9]. Solid stresses in tumors arise from two main sources: internal residual stresses generated by interactions among tumor cells and extracellular matrix components, including collagen fibers and glycosaminoglycans, and external stresses imposed by surrounding normal tissues that resist tumor expansion [10]. Together, these fluid- and solid-phase forces create a mechanically confined microenvironment that can compress tumor cells, alter cell volume, and regulate cancer cell behavior.

Volumetric compression is a universal phenomenon across various tissue types, arising from osmotic pressure during digestion, mechanical compression surrounding tusmors, and increased tissue stiffness during fibrosis. By altering cellular water content and biomacromolecular concentration, volumetric compression regulates cell function across molecular, mechanical, and systems levels, including molecular crowding, biomolecular crowing, cell mechanics, and regulatory network remodeling [11]. Volumetric compression can be modeled using osmolarity adjustment, or mechanical compression. Osmotic compression provides a useful approach to impose sustained volumetric stress on cultured cells. Specially, polyethylene glycol (PEG) - based osmotic treatment has been used to modulate extracellular osmolality and compressive stress, leading to changes in cell volume, proliferation, migration, and nuclear growth [12]. In PEG-mediated volumetric compression, extracellular osmolarity is increased using polyethylene glycol or related osmotic agents, which draws water out of cells and reduces cell volume. Previous studies have shown that PEG- or osmolarity-induced volumetric compression can regulate breast cancer cell size, proliferation, cell-cycle dynamics, migration, and recovery after stress release [13, 14]. Prior studies have shown that hyperostmotic stress can reduce proliferation and migration in both high metastatic MDA-MD-231 cells and weakly metastatic MCF-7 breast cancer cells, and that some responses to mild osmotic stress are reversible after stress release [15]. Other work has shown that volumetric compression can regulate breast cell cluster migration and that cell - cell adhesion influences the sensitivity of tumorigenic clusters to osmotic stress [16]. A recent study showed that volumetric compressive stress caused by ECM drives an invasive phenotype and apoptosis-resistant in liver cancer [17] and breast cancer [18]. These studies support PEG-mediated volumetric compression as a physiologically relevant platform to investigate how cancer cells adapt to sustained compression and whether prior compression creates a mechanical memory that promotes migration and invasion after recovery.

During tumor progression, migratory cancer cells move between mechanically distinct new niches may retain mechanosensitive adaption acquired from prior mechanical conditions, allowing previous physical cues to influence subsequent cell behaviors, including migration, invasion, and metastatic potential [7, 19–22]. Compressive stress may function as more than an acute physical constraint; it can also act as a memory-encoding mechanical cue that influences later tumor cell behavior. Recent work demonstrated that compressive stress in mammary tumors activates Piezo1-dependent Rho - ROCK mechanotransduction and induces persistent tumor-promoting epigenetic changes, supporting the concept of compression-driven mechanical memory in breast cancer progression [7]. Although mechanical memory has been increasingly investigated in the context of cancer progression, most studies focused on matrix stiffness-induced memory, how compression-induced memory is maintained and translated into migratory potential remains poorly understood [23, 24]. This gap is important because cancer cells within a growing tumor may experience sustained volumetric compression caused by ECM accumulation, tissue confinement, and elevated solid stress, followed by partial mechanical release during matrix degradation, invasion into adjacent tissues, or treatment-induced changes in tumor architecture. Therefore, if prior compression primes cancer cells to become more migratory and invasive after release, compressive stress should be viewed not only as physical constraint but also as a potential memory-encoding cue that promotes tumor dissemination. Therefore, studying the recovery phase after volumetric compression allows us to determine whether cancer cells retain a pro-invasive post-compression phenotype associated with F-actin remodeling, YAP-associated mechanotransduction, and enhanced invasion.

In this study, we established a PEG-mediated volumetric compression and recovery model to investigate how sustained compression and subsequent release regulate cancer cell phenotypic behaviors, including proliferation, migration, and invasion. We first evaluated the tolerance of sustained volumetric compression by monitoring daily morphological changes, including cell area, perimeter, circularity, and shape index. We next investigated the cell shape dynamics of cancer cells post-compression and results revealed several phenotypic behaviors during recovery phase. We further studied the mechanotranduction by evaluating YAP nuclear translocation by quantifying nuclear and cytoplasmatic YAP expression ratio. Lastly, we examined collective cell migration and 3D spheroids invasion. Our results show that sustained volumetric compression constrains MCF-7 cells into a jammed-like, low-motility state, whereas release from compression induces a distinct post-compression recovery phenotype characterized by cytoskeletal remodeling, altered YAP localization, enhanced wound closure, and increased spheroid invasion. By analyzing both the compression and recovery phases, this work identifies post-compression recovery as an underexplored mechanobiological state and suggests that volumetric compression may function not only as an acute physical constraint but also as a mechanical-history cue that primes breast cancer cells for enhanced motility and invasion.

## Material and Methods

### Cell culture and reagents

Breast cancer cells (MCF-7) were originally acquired from ATCC. Cells were cultured and maintained in complete culture medium - DMEM supplemented with 10% fetal bovine serum (FBS) and 1% penicillin–streptomycin with medium change every two days. Cells were maintained in a humified incubator with 5% CO2 at 37 °C.

### Volumetric compression in cell culture

Two different osmotic pressure-induced volumetric compression medium were prepared using filter-sterilized polyethylene glycol 200 (PEG200) at the concentration of 2% and 4% (vol/vol). The isotonic medium (control) was added 1XPBS at the concentration of 4% to match the slight dilution of the serum in the compression medium. The compression condition medium was prepared a day before the experiment to allow PEG full incorporation. For 2D experiments, cells were seeded in 12-well plate at the concentration of 1x10^5^ cells/well. Cells were cultured in regular medium until reached 60% confluency. The medium was then replaced with isotonic, 2% PEG, and 4% PEG medium and continued cultured for 4 days. Then the medium was changed to a complete culture medium for recovery.

### Tumor growth in the 3D niche

To form spheroids, MCF-7 cells were harvested and counted. 1,000 cells were seeded per well in 100 µL of complete medium in ultra-low-attachment 96-well U-bottom plates (Corning, #4515) and cultured for 4 days to allow spheroids to form. Spheroids were collected using wide-bore pipette tips, allowed to settle for 3 - 5 min, and the supernatant was removed. Type I collagen (6 mg/mL stock) was neutralized on ice to a final concentration of 2 mg/mL and gently mixed with spheroids (100 µL cell suspension + 50 µL collagen). Glass-bottom wells were pre-coated with polydopamine at a concentration of 1.8 mg/mL for overnight to promote gel anchorage.

Then the spheroid - collagen mixture (50 uL) were added to each well to form gel droplets. The droplets formed domes and were allowed to solidify for 1hr at 37°C, followed with culture medium. Spheroids embedded in collagen were imaged daily, with medium replacement every two days.

### Cell proliferation and viability

The cell proliferation and viability after PEG 200 incubation were evaluated using cell counting kit-8 (cck-8, Sigma Aldrich), following the manufacturers’ instructions. Cells were seeded at a concentration of 1000 cells per well in 96-tissue culture well plates with a volume of 100 μL culture medium. For the cell toxicity test, cells were treated with PEG at different concentrations (isotonic, 2%, and 4% PEG-200) after 1 day of incubation. For cell proliferation assay, cells were incubated with PEG-200, and cell proliferation was evaluated at different time points, 4 hr, 8 hr, 12 hr, and 24 hr. After incubation, cck-8 was added to the cells and cultured for 4 h. The absorbance was measured at 450 nm for each sample and compared using a fluorescence microplate reader (BioTek, Synergy 2).

### Live/dead viability staining

The cell viability was further evaluated using live/dead viability assay (ThermoFisher). Cells were stained using propidium iodide (PI, 10 μg/mL), a fluorescent agent that binds to DNA by intercalating between the bases with little or no sequence preference. Hoechst 33342 was used to stain the cell nucleus (1:2000) for 30 minutes. After staining, cells were washed three times with 1× PBS to remove the extra dye. Cells were then imaged using Texas Red (535/617 nm) and DAPI (360/460 nm) filters using an Echo Revolution fluorescence microscope.

### Immunofluorescence staining

For YAP immunostaining, cells were fixed with 4% paraformaldehyde for 15 min, permeabilized with 0.1% Triton X-100 for 10 min, and blocked with 5% bovine serum albumin for 1 h at room temperature. Samples were incubated with a primary antibody against YAP (Santa Cruz, sc-101199, 1:200) overnight at 4°C, next day, cells were washed 3 times using 1x PBS, and cells were incubated with Alexa Fluor 488 Goat anti-Mouse secondary antibody (ThermoFisher) 1:1000 for 2 h at room temperature. For F-actin staining, cells were incubated with phalloidin (1:30) and Hoechst 33342 (1:2,000) for 30 min at room temperature. The cells were then washed 3 times with ×1 PBS before fluorescence imaging and analysis. Cell nuclei were counterstained with Hoechst 33342, and fluorescence images were acquired using identical imaging settings across all experimental groups. YAP nuclear translocation was quantified by calculating the ratio of nuclear to cytoplasmic YAP fluorescence intensity using the equation: 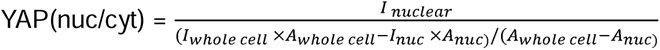, where *I* indicates average fluorescence intensity.

### Wound scratch assay

Cell migration was assessed using a scratch assay. MCF-7 cells were seeded in 12-well plates at a density of 40,000 cells per well in 700 µL DMEM and cultured until a confluent monolayer formed. A straight scratch was created across the cell layer using a 200-µL pipette tip, ensuring a consistent wound width. Detached cells and debris were removed by washing, and wells were replenished with fresh DMEM. Migration of cells into the wound area was then monitored by live-cell imaging, with images acquired at 30-minute intervals to quantify wound closure dynamics.

### Imaging and statistical analysis

Images were obtained using an Echo Revolution fluorescence microscope with an integrated digital camera (5MP CMOS Color for bright field, 5 MP sCMOS Mono for fluorescence imaging).

To ensure consistency, all images were captured under identical settings, including exposure time and gain. Image analysis and data collection were conducted using NIH ImageJ software. Statistically significant differences between the means of two groups were assessed using a Student’s *t* test, whereas data containing more than two experimental groups were analyzed with a one-way ANOVA test. Error bars in all figures represent SED. All statistical analyses were performed in the Origin 9.0 software. \**P* < 0.05, \*\**P* < 0.01, and \*\*\**P* < 0.001, respectively.

## Results

### Effects of volumetric compression on cell viability

To evaluate the cellular response to PEG 200-mediated volumetric compression, MCF-7 cells were cultured under isotonic conditions or exposed to 2% or 4% PEG and assessed using propidium iodide (PI) staining assay. Cells maintained in isotonic medium showed normal morphology and minimal PI signal, indicating preserved cell viability. Exposure to PEG produced concentration-dependent changes in cell morphology and viability-associated staining, with 4% PEG causing more pronounced cell rounding, reduced cell density, and increased PI-positive signal compared with 2% PEG. Quantitative analysis further showed a time- and concentration-dependent decrease in normalized cellular intensity under PEG treatment, with 2% PEG producing a moderate reduction and 4% PEG producing a stronger decrease, particularly after 24 h. We selected 4, 8, 10, and 24 h to capture early, intermediate, and late acute responses to PEG 200-mediated volumetric compression, allowing us to monitor the onset, progression, and persistence of morphology and viability changes before the longer-term compression and recovery experiments. These results indicate that PEG-mediated volumetric compression affects MCF-7 cell morphology and viability in a dose-dependent manner, supporting the use of 2% PEG as a compression condition that induces measurable cellular stress while maintaining sufficient cell survival for subsequent recovery, migration, and invasion analyses.

### Recovery from volumetric compression promotes cell spreading and shape remodeling

To investigate phenotypic behavior of cancer cell recovery process after sustained compression, cancer cells were exposed to PEG 200 with the concentration of 2% and 4% for four days, and then the volumetric compression was removed, and cells were replaced with fresh DMEM for the recovery period. Cell morphology was tracked daily to monitor the cellular behaviors under volumetric compression and during recovery process. **Figure 2A** shows the timeline of the experimental design. Here we chose four days of sustained compression to make sure cells can recover and reach the highest tolerance of compression. Longer than 4 days of compression cause majority of cell death due to the compression. For 4% sustained compression and recovery, the results showed no recovery after compression removal. All cells suffered and experience death even after compression releases. Thus, we focused on 2% PEG induced volumetric compression. **Figure 2B** shows the daily tracking of bright field images of MCF-7 at the same location for cells under compression for 4 days and recovery process for another 3 days. It is evident that MCF-7 cells experience size reduction with reduced area and perimeter due to the water drawn caused by the volumetric compression. Representative daily images and morphometric tracking show that MCF-7 cells undergo remarkable morphological remodeling during the compression - recovery timeline. During the early compression phase, MCF-7 under compression shown more confined, compact, and less dynamically remodeled, with reduced cell spreading and limited expansion of the cell-covered area. MCF-7 under compression also shows less cell displacement and fewer visible protrusion, suggesting that volumetric compression restricted cell rearrangement and suppressed morphological plasticity. After release from PEG 200-mediated volumetric compression, however, recovered cells progressively increased in size, as shown by the expansion of representative cell outlines from day 5 to day 8. Quantitative analysis, **Figure 2C-2F**, confirmed this recovery - associated remodeling, with both cell area and perimeter increasing substantially over time, indicating enhanced spreading and cell boundary expansion after compression release. In parallel, aspect ratio gradually decreased while roundness increased during the recovery period, suggesting that recovered cells transitioned from a more elongated or constrained morphology toward a larger, more isotropically spread phenotype. Together, these results indicate that post-compression recovery is associated with pronounced cell shape remodeling, supporting that MCF-7 cells do not simply return to their original morphology after volumetric compression but instead acquire a distinct recovery-associated morphodynamic state.

**Figure 1.**
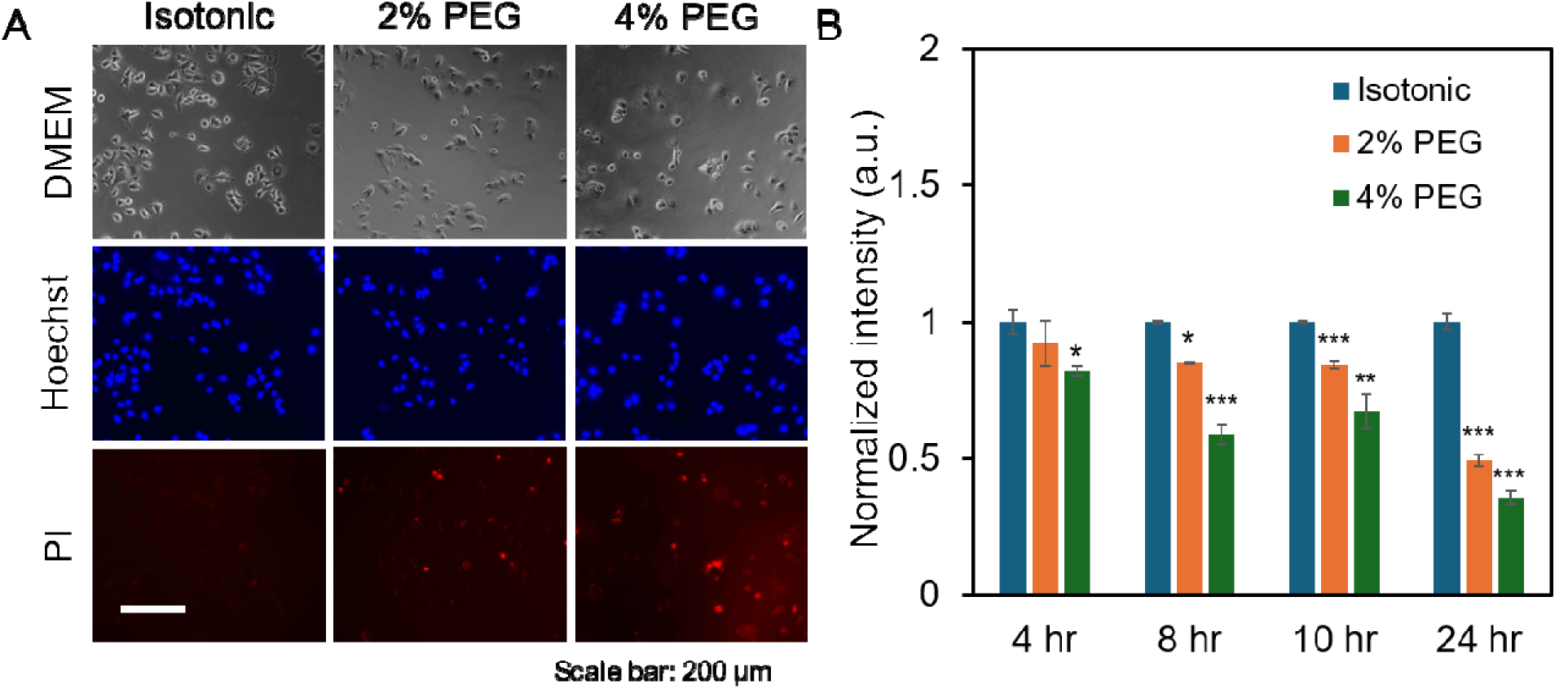
PEG-induced osmotic compression reduces MCF-7 cell viability in a concentration- and time-dependent manner. (A) Representative phase-contrast images (top), nuclear fluorescence images (blue; middle), and dead-cell fluorescence images (red; bottom) of MCF-7 cells cultured under isotonic, 2% PEG, and 4% PEG conditions, shown from left to right. Scale bar: 200 µm. (B) Relative cell viability after 4, 8, 10, and 24 h of treatment, normalized to the corresponding isotonic control. Data represents over 100 cells in each group and are expressed as mean ± s.e.m. A two-tailed t-test was used to analyze differences between control and shear conditions. (n = 5, ***, p < 0.001, **, p < 0.01, *, p < 0.05).

**Figure 2.**
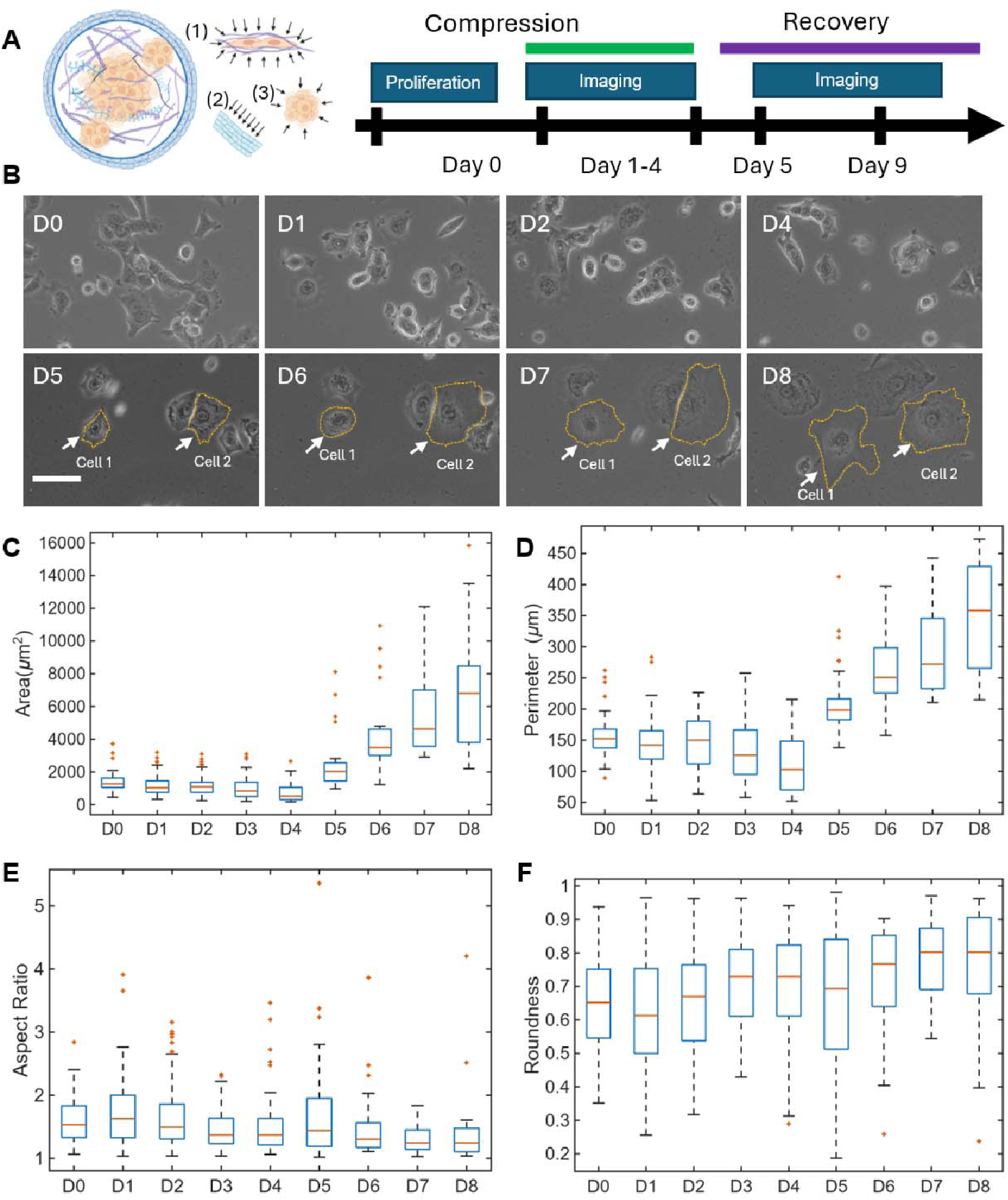
Daily morphological tracking of MCF-7 cells under volumetric compressive stress. (A) Left: Tumor expansion in a confined environment can lead to: (1) mechanical compression on the tumor edge cells against a stiff, fibrotic substrate; (2) compression on the adjacent normal cells; and (3) volumetric compression on the cells toward the core of the tumor. Right: Experimental design. (B) Representative images of cells under compressive stress after different days of compression. Scale bar: 100 µm. (C-F) Comparison of area, perimeter, aspect ratio and roundness of cells at different time points.

### Post-compression recovery drives progressive cell motility and morphological remodeling

To investigate whether MCF-7 cells retain a mechanical memory after PEG – mediated volumetric compression, we performed single-cell time – lapse tracking after recovery period and compared their behavior with control group. Representative time-lapse bright field images showed that cells maintained relatively compact morphologies with modest boundary remodeling over 14 h for control group, whereas recovered cells displayed more pronounced shape changes, including dynamic protrusions, spreading, and retraction, **Figure 3A**. To better understand whether prior compression alters subsequent cell motility, we quantified cell trajectories, cell area, perimeter, aspect ratio, circularity, and shape index over time (**Figure 3C-3H, Figure S1**). Cell trajectory analysis further demonstrated that post-compression recovered cells exhibited altered migration paths compared with control cells, indicating that prior compression history changes subsequent cell movement behavior after PEG removal, **Figure 3C-3D**. Quantitative tracking of cell area showed that recovered cells occupied a substantially larger area than control cells, consistent with enhanced spreading and cytoskeletal remodeling during recovery **Figure 3E-3F**. In parallel, time-dependent analysis of shape index revealed dynamic fluctuations in cell shape, reflecting continuous remodeling of the cell boundary during migration **Figure 3G-3H**. Together, these single-cell tracking results indicate that recovery from volumetric compression produces a distinct post-compression phenotype in which MCF-7 cells show altered trajectories, increased spreading, and sustained morphodynamic activity compared with control group.

**Figure 3.**
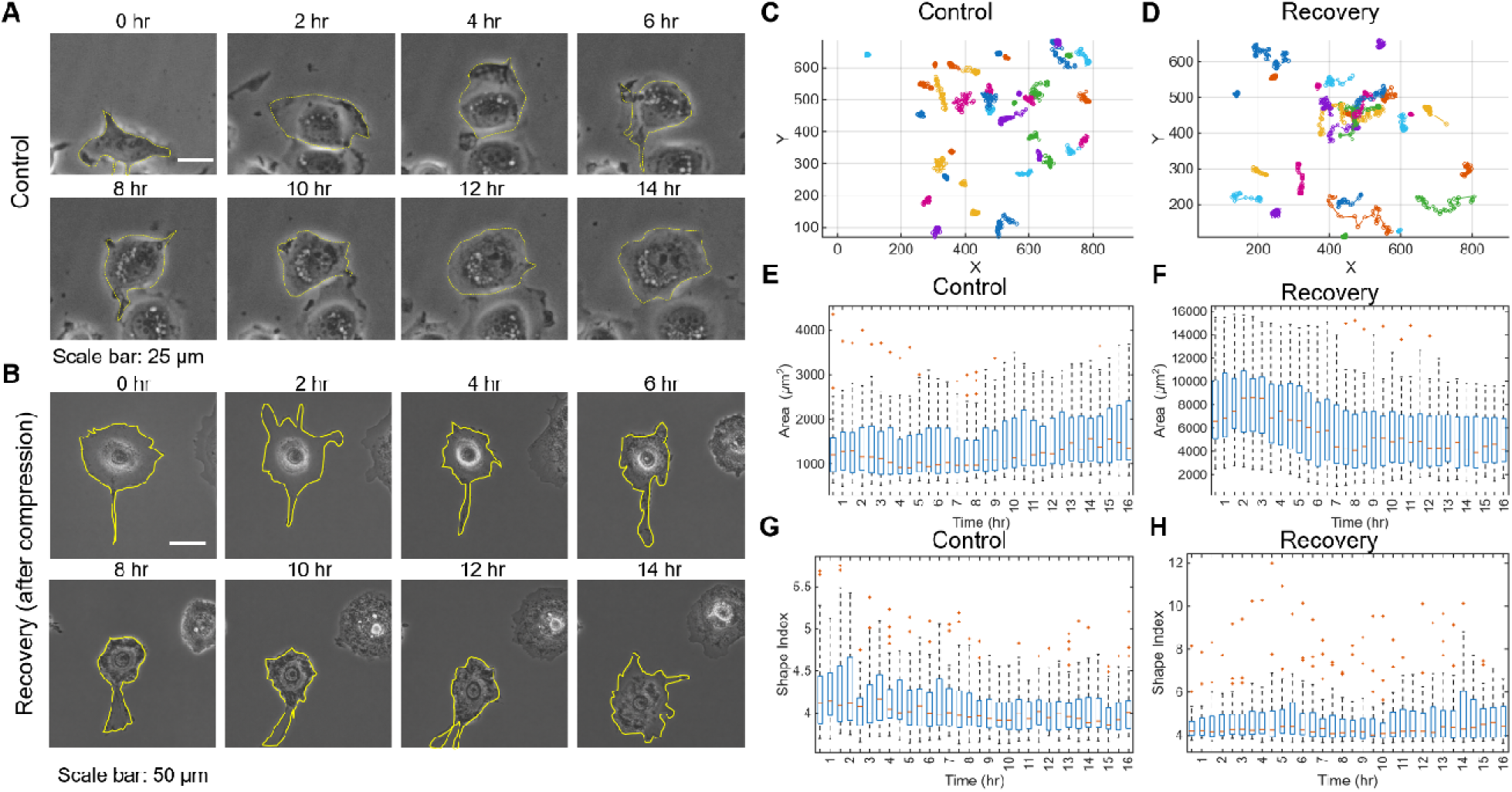
Quantitative analysis of dynamic cell morphology tracking for isotonic and recovery cells. (A-B) Representative images of cells without compression and recovery cells. Scale bar: 50 µm. (C-D) Trajectory of 35 cells for control and recovery group. (E-H). Quantitative analysis of cell area and shape index tracking over time. Data represents over 40 cells in each group.

### Post-compression recovery enhanced YAP nuclear translocation and cytoskeletal remodeling

During cell recovery process, we noticed several cell behaviors that are worth noting. Cells experiences large expansion with 10 times of increase of cell area (giant single nucleated cell, GSNCs), or giant muclinucleated cells (GMNCs), and regular-sized recovery cells (RCs). Thus, we categorized the recovery cells into three group: GSNCs, GMNCs, and RCs. We examined phenotypic behaviors and mechanotransduction by measuring YAP nuclear translocation. As shown in **Figure 4A**, F-actin staining revealed distinct cytoskeletal remodeling patterns among control cells, GSNCs, GMNCs, and RCs following PEG 200-mediated volumetric compression and recovery. Because volumetric compression can regulate cellular function by altering molecular crowding, biomolecular condensate formation, cell mechanics, and systems-level regulatory networks, we reasoned that post-compression recovery may generate heterogeneous cytoskeletal states. Compared with control cells, GSNCs and GMNCs exhibited markedly enlarged cell bodies and prominent F-actin remodeling. GSNCs showed strong peripheral F-actin localization, suggesting an early recovery-associated response in which cortical actin reorganizes during cell re-expansion after compression release. GMNCs displayed F-actin signal across both the cell cortex and cytoplasmic region, indicating a more advanced or active remodeling state with broader cytoskeletal redistribution. In contrast, RCs showed a more compact, ring-like F-actin organization that more closely resembled the control phenotype. These observations indicate that PEG 200-mediated volumetric compression and recovery do not produce a uniform cellular response; instead, they generate heterogeneous post-compression phenotypes with distinct F-actin architectures that may reflect different stages of cytoskeletal remodeling and mechanical adaptation. We next quantified YAP nuclear-to-cytoplasmic (nec/cytop) ratio using mean fluorescence intensity by calculating the average pixel intensity within the nuclear region divided by the average pixel intensity in the cytoplasmic region. For GMNCs, average nuclear intensity were measure by calculating all the nuclears within the same cells. Our results revealed that GSNCs exhibited the highest YAP nuc/cyto ratio, followed by GMNCs and RCs, whereas control cells showed the lowest YAP nuc/cyto ratio. This finding suggests that volumetric compression and recovery activate YAP -associated mechanotransduction, particularly in enlarged post-compression cell states. The stronger YAP nuclear localization in GSNCs may reflect an early recovery response associated with rapid cell -re-expansion and cortical cytoskeletal remodeling, while the intermediate YAP levels in GMNCs and RCs suggest progressive remodeling toward more stabilized recovery states. Quantitative analysis of nuclear morphology further indicated that compression and recovery affected nuclear size, shape, or circularity, consistent with the idea that osmotic compression influences both the cytoskeleton and nucleus. Together, these results suggest that post-compression recovery is not merely a passive relaxation process but is associated with coordinated F-actin reorganization, altered cell and nuclear morphology, and YAP-associated mechanotransduction. These cytoskeletal and nuclear changes provide a possible mechanobiological basis for the enhanced migration observed in recovered MCF-7 cells.

**Figure 4.**
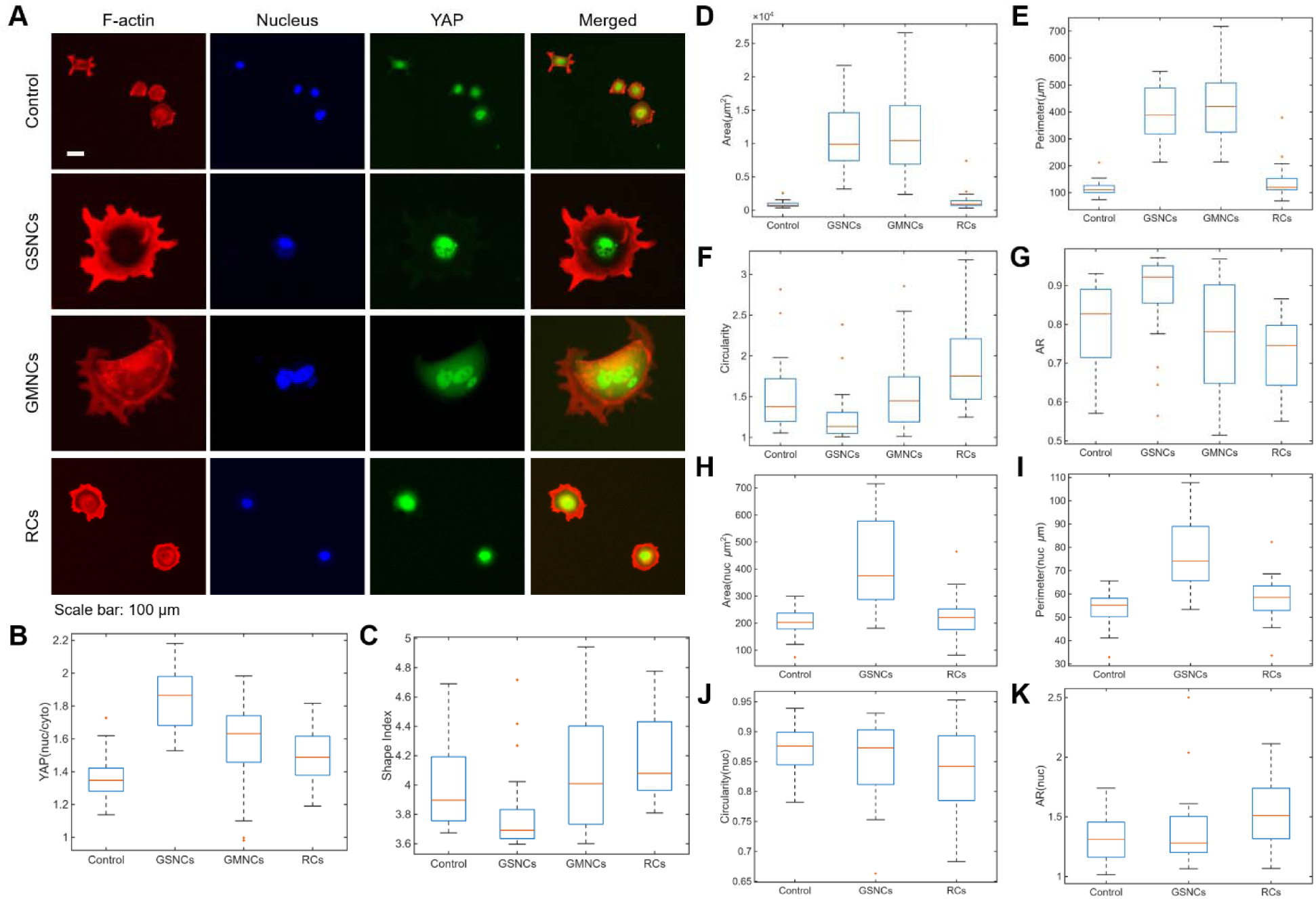
F-actin organization, YAP localization, and cell and nuclear morphology following PEG-induced osmotic compression and recovery. (A) Representative fluorescence images showing F-actin (red), nuclei (blue), YAP (green), and merged channels in control cells, giant single-nucleated cells (GSNCs), giant multinucleated cells (GMNCs), and RCs. Scale bar, 100 μm. (B) Quantification of the nuclear-to-cytoplasmic YAP fluorescence-intensity ratio. (C) Cell shape index. (D–G) Quantification of whole-cell area, perimeter, circularity, and aspect ratio (AR), respectively, in control cells, GSNCs, GMNCs, and RCs. (H–K Quantification of nuclear area, perimeter, circularity, and AR, respectively, in control cells, GSNCs, and RCs. GMNCs were excluded from the single-nucleus morphological analysis because individual cells contained multiple nuclei. Box plots show the median, interquartile range, whiskers, and individual outliers. Data represents over 40 cells in each group. (n=6)

### Post-compression recovery increased wound closure rate

To determine whether compression induced morphological and cytoskeletal remodeling translate into functional changes in collective cell migration, we performed wound healing assays under isotonic, compressed, and post-compression recovery conditions. **Figure 5A** show representative images of collective migration at different time points, including 0hr, 4 hr, 8 hr, 12 hr, and 16 hr. For control group, MCF-7 cells progressively migrate into the wound area, resulting in gradual wound closure over time. In contrast, cells under PEG-mediated volumetric compression showed markedly impaired collective migration, with limited movement into the wound gap and delayed wound closure. Notably, cells exposed to four days of PEG 200-mediated volumetric compression followed by four days of recovery exhibited accelerated wound closure compared with both compressed and control groups. Specifically, **Figure 5B** shows that PEG -mediated volumetric compression strongly suppresses MCF-7 wound closure, while post-compression recovery enhances collective migration. The compressed group remained nearly unchanged throughout the 16 h imaging period, indicating that cells under compression were largely jammed or migration-arrested. In contrast, control cells gradually closed the wound over time, reaching approximately 20–25% closure by 16 h. Recovered cells showed a faster and greater wound closure response than control cells, reaching approximately 30–35% closure by 16 h, suggesting that prior compression followed by recovery primes MCF-7 cells for enhanced collective migration. **Figure 5C** show the migration rate comparison at different time points. Control cells exhibited relatively modest and stable migration rates over time, whereas recovered cells showed a progressive increase in migration rate, especially at later time points. Together, these two panels demonstrate a biphasic response: active volumetric compression suppresses MCF-7 migration, whereas post-compression recovery induces a enhanced migratory with faster wound closure and increased migration rate.

**Figure 5.**
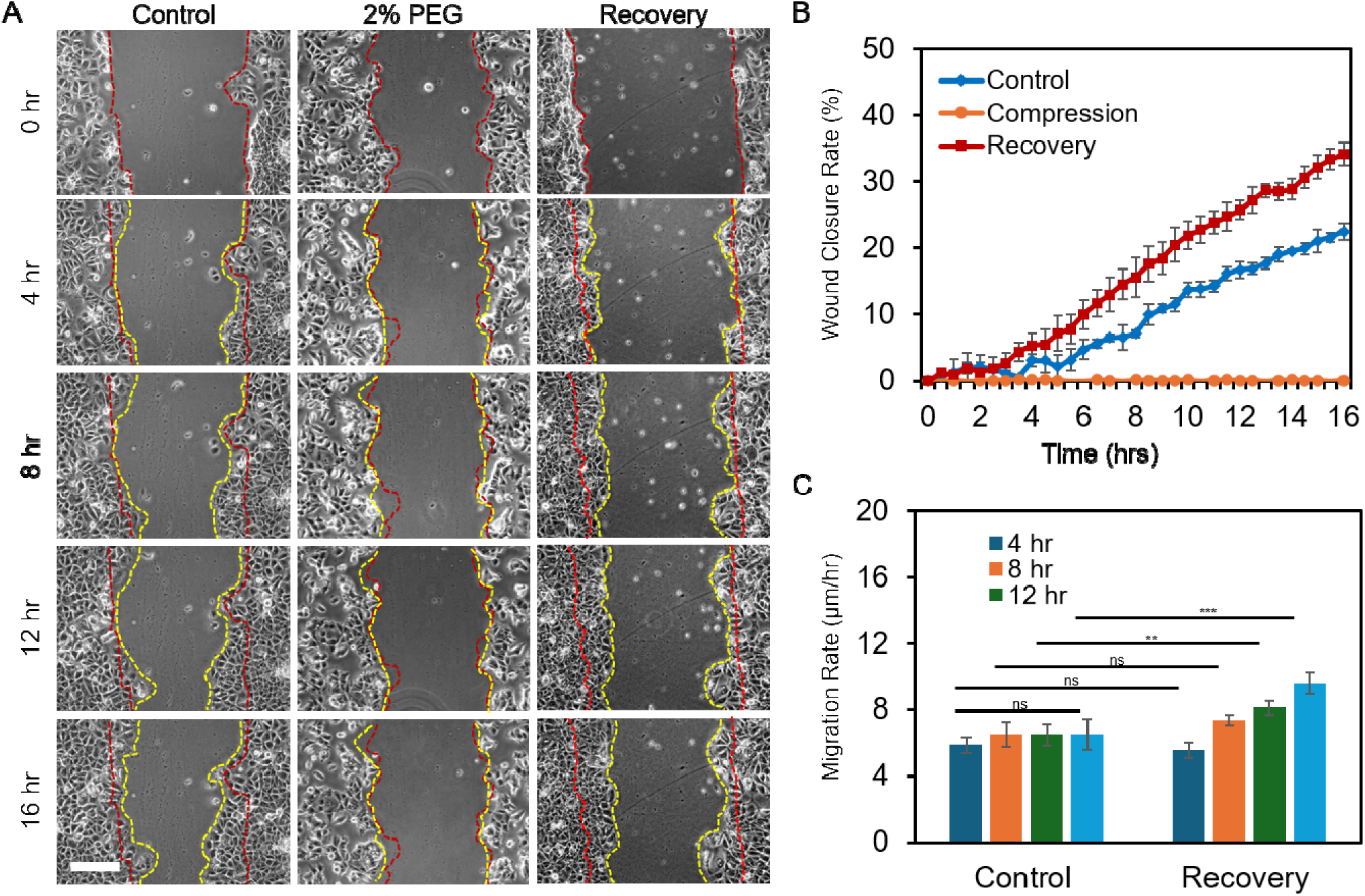
PEG-induced osmotic compression suppresses MCF-7 collective cell migration, whereas post-compression recovery enhances wound closure. (A) Representative phase-contrast images of scratch wounds in control, 2% PEG-compressed, and recovered MCF-7 cell monolayers at 0, 4, 8, 12, and 16 h. Red dashed lines indicate the initial wound boundaries at 0 h, whereas yellow dashed lines indicate the wound edges at the corresponding subsequent time points. Scale bar, [XX μm]. (B) Quantification of wound closure over 16 h. Wound closure was substantially inhibited during 2% PEG treatment, whereas recovered cells exhibited faster closure than control cells. (C) Migration rates of control and recovered cells calculated at 4, 8, 12, and 16 h. Data represents over 100 cells in each group and are expressed as mean ± s.e.m. A two-tailed t-test was used to analyze differences between control and shear conditions. (n = 5, ***, p < 0.001, **, p < 0.01, *, p < 0.05).

### Post-compression recovery enhances MCF-7 spheroid invasion and cell dissemination

To determine whether post-compression recovery enhances invasive behavior in a 3D tumor-like context, MCF-7 spheroids were monitored for four days under isotonic, compressed, and recovery conditions. Representative images showed that isotonic spheroids exhibited gradual expansion with limited cell dissemination, whereas spheroids maintained under PEG 200-mediated volumetric compression remained relatively compact, with minimal invasive outgrowth, **Figure 6A**. In contrast, recovered spheroids displayed a pronounced increase in radial invasion and cell dissemination over time, with visible release of individual cells from the spheroid boundary by days 3 - 4. Quantitative analysis confirmed this phenotype, showing that recovered spheroids had substantially greater invasion length and a higher number of isolated disseminated cells compared with isotonic controls, particularly at day 4, **Figure 6B-6C**. These results indicate that sustained volumetric compression suppresses spheroid invasion during active compression, but release from compression promotes a strong pro-invasive rebound, supporting the concept that prior compression can generate a post-compression mechanical memory that enhances MCF-7 spheroid invasion and cell dissemination.

**Figure 6.**
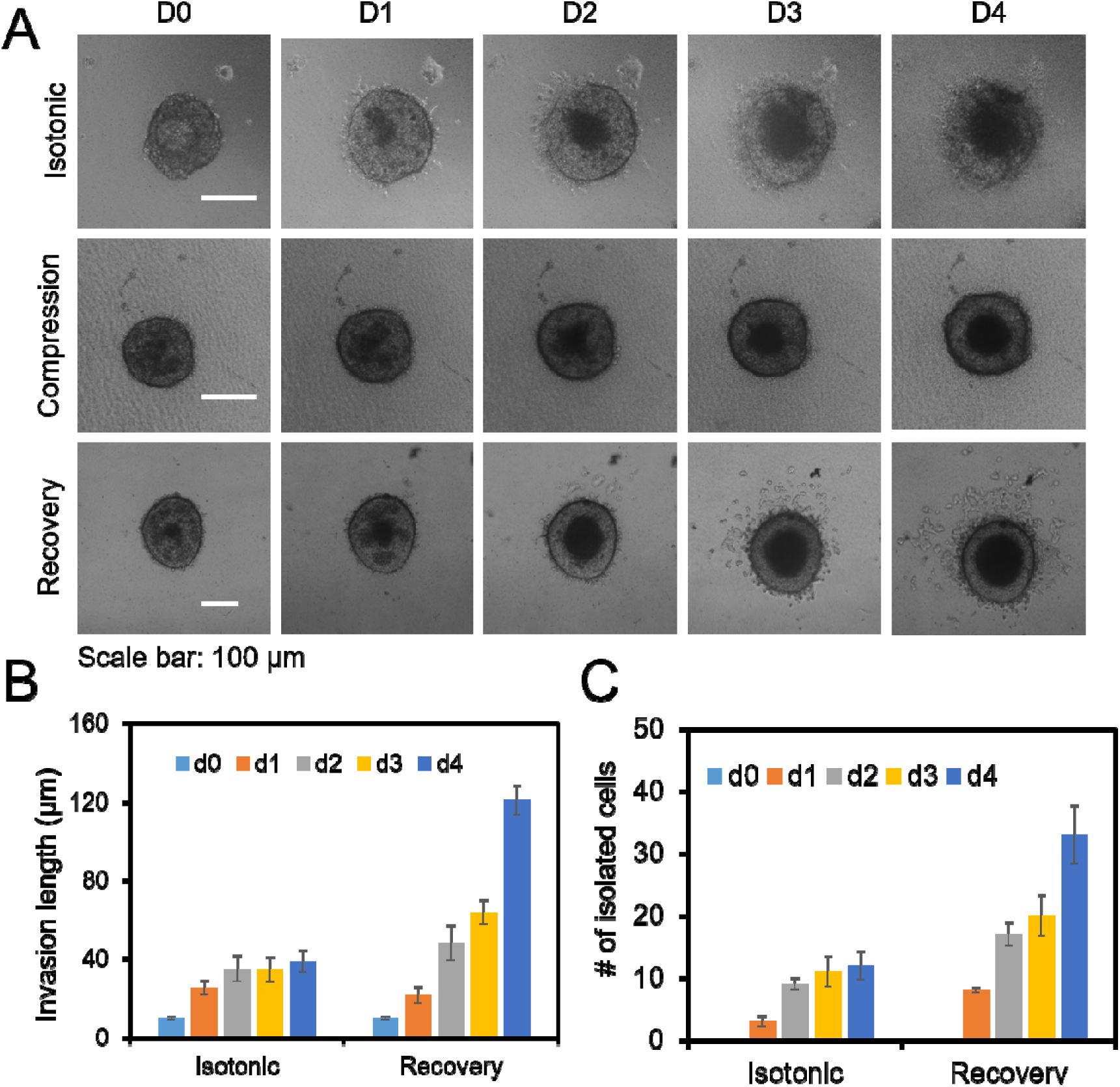
Recovery from PEG-induced osmotic compression enhances MCF-7 spheroid invasion and cell dissemination. (A) Representative phase-contrast images of MCF-7 spheroids maintained under isotonic conditions, subjected to PEG-induced osmotic compression, or allowed to recover following compression. Spheroid morphology and invasion were monitored from day 0 (D0) to day 4 (D4). Scale bars, 100 μm. (B) Quantification of invasion length in isotonic and recovered spheroids over the 4-day observation period. (C) Quantification of isolated cells that detached from and migrated away from isotonic and recovered spheroids. Data represents over 20 spheroids in each group and are expressed as mean ± s.e.m. A two-tailed t-test was used to analyze differences between control and shear conditions. (n = 5, ***, p < 0.001, **, p < 0.01, *, p < 0.05).

## Discussion

In this study, we established PEG-mediated volumetric compression and recovery model to investigate how sustained compression history regulate MCF-7 breast cancer cell morphology, cytoskeletal organization, YAP nuclear localization, collective cell migration and 3D spheroid invasion. Our results revealed that during active compressive, MCF-7 cells exhibited reduced cell area and perimeter, limited morphodynamic remodeling, suppressed collective migration, and reduced spheroids invasion in 3D collagen gel. In contrast, during recovery, cells displayed increased area and perimeter, altered single-cell trajectories, enhanced morphodynamic fluctuations, distinct F-actin remodeling, increased YAP nuclear/cytoplasmic localization in specific post-compression cell states, accelerated wound closure, and increased 3D spheroid invasion. Together, these findings support the concept that sustained volumetric compression transiently constrains MCF-7 cells into a jammed-like, low-motility state, whereas post-compression recovery generates a mechanically primed phenotype with enhanced migratory and invasive capacity.

Volumetric compression plays a critical role in cancer progression by acting as a mechanical driving force that alters gene expression, promotes drug resistance, and changes in cell behavior within the tumor microenvironment [25, 26]. Volumetric compression could alter intracellular calcium signaling and activated survival pathways (such as those mediated by Rac1 and YAP), helping cancer cells resist programmed cells death [27–29]. Previous studies also show compression could increase cell heterogeneity, where reducing a cell’ s volume via physical crowding drive diverse and aggressive cell fates, such as epithelial – mesenchymal transition (EMT) linked to invasion [27]. Our findings are consistent with the concept that volumetric compression is not merely a passive reduction in cell size, but a multiscale mechanobiological input that regulates cell function through changes in water content, biomacromolecular concentration, condensation, and molecular crowding. Volumetric compression has been described as a pervasive biological process that occurs during tumorigenesis and can influence cellular function by modulating biochemical reaction kinetics and equilibria. In addition, cell volume is coupled to cell mechanics, molecular crowding, and biomolecular concentrations, allowing extracellular physical cues to regulate intracellular biochemical and genetic processes [11]. In our study, PEG 200 increases extracellular osmolarity, which drives water efflux and reduces cell volume; this is consistent with the principle that increased extracellular osmotic pressure removes water from cells and produces a more concentrated intracellular environment [11]. Prior studies have also shown that water efflux and volumetric compression can alter cell stiffness and fate decisions [30], while compression-induced intracellular crowding can regulate signaling pathways such as Wnt/β-catenin [12] . Therefore, the morphological and cytoskeletal changes observed in our study likely reflect both direct physical effects of volume reduction and downstream adaptive remodeling after compression release.

It is worth mentioning that the emergence of giant multinucleated cells after PEG -mediated volumetric compression and recovery may represent a stress-adapted intermediate state linking compression history to enhanced migration and invasion. Polyploid giant cancer cells (PGCCs), including multinucleated giant cancer cells, are associated with tumor heterogeneity, treatment resistance, recurrence, and metastasis, and they can arise through several mechanisms, including cell fusion, cytokinesis failure, mitotic slippage, endoreplication, and cell cannibalism. Notably, cell fusion can generate hybrid cells with chromosomal instability and enhanced malignant traits, including increased growth, spread, drug resistance, stem-like features, and tumor diversity [31–33]. In our study, the enlarged multinucleated cells observed after recovery may therefore reflect a PGCC-like stress-adapted phenotype induced by volumetric compression, which is consistent with the broader concept that volumetric compression alters cell mechanics, molecular crowding, and systems-level regulatory networks. Importantly, daughter cells derived from PGCCs have been reported to acquire enhanced invasion, metastasis, and therapy resistance, and cytoskeletal remodeling has been implicated in promoting migration and invasion of PGCCs and their progeny. Thus, giant multinucleated cells formed during post-compression recovery may serve as a transient mechanically adapted state that contributes to the enhanced collective migration and spheroid invasion observed at later time points.

The suppression of MCF-7 motility during active volumetric compression supports the idea that healing assays, compressed cells showed minimal movement into the wound gap, while compressed spheroids remained compact with limited invasive outgrowth. This behavior is consistent with the broader jamming, in which epithelial or cancer cell collectives can transition between solid-like, low-motility states and fluid-like, migratory states depending on cell shape, adhesion, and mechanical constraints [34]. Park et al. established cell shape and collective motion as key features of epithelial unjamming, and Cai et al. later showed that compressive stress can drive adhesion-dependent jamming or unjamming transitions in breast epithelial and breast cancer cells [34, 35]. However, our data extends this by emphasizing the recovery phase after sustained volumetric compression. While previous studies have shown that mechanical compression can promote invasive phenotypes in cancer cells during active loading [18], our results indicate that the strongest pro-migratory phenotype emerges after compression is removed. This distinction suggests that compression history, rather than compression alone, may be a critical regulator of subsequent breast cancer cell behavior.

F-actin, YAP, and spheroid invasion data suggest that PEG 200-mediated volumetric compression creates a recovery-associated mechanobiological state rather than a simple return to baseline. Recovered cells displayed heterogeneous F-actin remodeling, including peripheral actin enrichment and broader cytoplasmic redistribution, consistent with volumetric compression altering molecular crowding, biomolecular condensates, and cell mechanics. YAP nuclear enrichment in enlarged recovered cells further suggests activation of mechanotransduction during recovery. Functionally, recovered spheroids showed greater invasion length and cell dissemination, indicating that release from compression permits mechanically primed cells to express enhanced invasive behavior in 3D. These findings suggest that active compression suppresses invasion, whereas release from sustained compression allows mechanically primed cells to express a more invasive phenotype, consistent with prior 3D breast cancer studies linking cell-volume regulation, swelling, softening, and invasion [13, 14]

A central implication of this work is that post-compression recovery may represent a form of mechanical memory. Mechanical memory describes the ability of cells to retain information from prior mechanical environments and use that information to regulate later behavior. This concept has been studied most extensively in the context of matrix stiffness, where prior exposure to stiff environments can induce persistent tumorigenic phenotypes through changes in chromatin accessibility [36]. More recent work has also shown that force-induced changes in nuclear and chromatin-associated protein dynamics can persist after force cessation, supporting the idea that cells can store mechanical history at the nuclear level [21]. Our results suggest that volumetric compression may also function as a memory-encoding cue. In breast tumors, cells may experience sustained confinement caused by tumor expansion, ECM accumulation, stromal resistance, and elevated solid stress, followed by partial mechanical release during ECM degradation, tissue invasion, vascular remodeling, or therapeutic remodeling of the tumor microenvironment [37, 38]. If prior compression primes cells for enhanced migration and invasion after release, then compression should be viewed not only as an acute physical barrier but also as a potential driver of future invasive behavior.

Future work will focus on identification of molecular and biophysical mechanisms underlying post-compression memory. Perturbation studies targeting F-actin remodeling, actomyosin contractility, YAP/TAZ, RhoA/ROCK, Piezo channels, and osmolyte transport will determine which pathways drive enhanced migration and invasion after recovery. Single-cell transcriptomic or epigenetic profiling should test whether recovered cells acquire stable states or heterogeneous subpopulations, as volumetric compression can induce cancer cell heterogeneity [39, 40]. These studies should be extended to invasive breast cancer lines, nonmalignant epithelial cells, patient-derived organoids, and ECM-rich 3D models.

## Data availability

The data supporting this article have been included as part of the ESI.**†**

## Acknowledgements

S. Wang acknowledges the financial support from NSF (2342274).

