## Supplemental Figure S1 for "Sustained Volumetric Compression Induces Cell Jamming and Primes Breast Cancer Cells for Enhanced Post-Compression Migration and Invasion"

**Supplemental Figures**

**Figure S1. Shape index change of MCF-7 cells post compression.**

Supplemental Figures

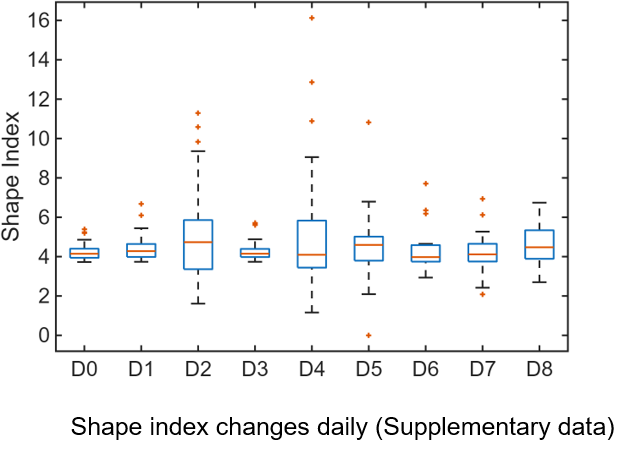

**Figure S1. Shape index change of MCF-7 cells post compression.**
